# Natural variation in *DXR* expression is associated with apical hook bending under NaCl stress in *Arabidopsis thaliana*

**DOI:** 10.64898/2026.04.12.718021

**Authors:** Elizabeth van Veen, Tirza van den Dikkenberg, René Boesten, Xizheng Chen, Joram A. Dongus, Charlotte M. M. Gommers

## Abstract

Successful soil emergence requires apical hook establishment during skotomorphogenesis. Soil salinity disrupts this process, but the mechanisms linking environmental stress to hook growth remain unclear. Here, we employed a genome-wide association study (GWAS) across *Arabidopsis thaliana* accessions to identify a locus associated with variation in the salt-induced reduction of apical hook curvature. Fine mapping reveals genetic variation in the promoter of *1-DEOXY-D-XYLULOSE 5-PHOSPHATE REDUCTOISOMERASE* (*DXR*), encoding the first committed enzyme of the plastid-localized methylerythritol phosphate (MEP) pathway. Accessions carrying an alternative promoter haplotype exhibit elevated *DXR* expression and a stronger hook bending under salt. Salt treatment and loss of transcriptional repressor PHYTOCHROME INTERACTING FACTOR1 (PIF1) additively increase *DXR* transcript levels, and *pif1*-2 seedlings exhibit higher apical hook angles under salt stress. This phenotype is suppressed by inhibition of DXR activity, indicating that an increased MEP pathway flux reduces salt sensitivity in *pif1-2* seedlings. Across accessions, *DXR* expression positively correlates with hook curvature under salt stress, further strengthening the link between DXR and modulation of hook bending under salt stress. Together, these findings identify plastid metabolism as a regulatory layer linking environmental stress to altered skotmomorphogenis, and raise important questions about how etioplast-derived signals interact with growth-control networks in the dark.

**Highlight:** A genome wide association study in Arabidopsis reveals that variation in *DXR* expression, encoding an essential enzyme in the MEP pathway, drives salt-induced repression of apical hook formation darkness.

## Introduction

After release from the mother plant, seeds can become buried in a soil environment, where they are protected from adverse weather conditions and predation. When conditions become favourable, germination will occur, and the young seedling emerges from the testa (1, 2). During this time, the seedling enters a heterotrophic growth program known as skotomorphogenesis, or etiolation, where development is directed toward emergence from the soil (3, 4). To achieve this, seedlings prioritize rapid elongation of the hypocotyl, as well as bending of the upper portion to form an apical hook, whilst restricting the expansion and opening of the pre-photosynthetic cotyledons (3-5). The apical hook is a crucial feature that protects against mechanical damage and allows smooth soil penetration, ensuring successful seedling establishment (6).

Etiolated growth is tightly regulated by the local environment, with (the absence of) light being the primary driver (3-5). Apical hook development relies not only on darkness to be formed (7), but is also impacted by mechanical pressure imposed by the soil (6, 8), oxygen availability (9), gravity (10), and temperature (11). In *Arabidopsis thaliana*, multiple phytohormone and environmental signalling pathways are integrated by two sets of transcription factors, ETHYLENE INSENSITIVE 3/EIN3-LIKE 1 (EIN3/EIL1) and the PHYTOCHROME INTERACTING FACTORs (PIFs; PIF1, PIF3, PIF4 and PIF5), to regulate apical hook development (12, 13). These transcription factors are best studied in the context of response to mechanical contact from the soil and light, respectively (7, 13, 14).

Soil salinization is a major environmental factor that constrains plant growth and development, affecting over 950 million hectares of land globally (15-17). Our earlier identification of salt-induced reductions in apical hook bending reveals a previously unrecognized detrimental effect of salinity on early seedling development, as salt-treated seedlings (similar to *ein3/eil1* mutants) exhibit impaired soil emergence (8, 18). We showed that salt stress disrupted the COP1 gradient in the apical hook, suggesting the activity of COP1, and transcription factors targeted by it, may act in a cell specific manner. Nevertheless, the mechanisms by which young seedlings sense salinity and transduce these signals towards the open hook phenotype remain unknown.

One way that seedlings adjust growth to fluctuating environmental conditions is by dynamically fine-tuning plastid metabolism (19, 20). A central component of this coordination is the methylerythritol phosphate (MEP) pathway, which serves as a sophisticated oxidative stress sensor by modulating levels of the intermediate metabolite methylerythritol cyclodiphosphate (MEcPP) (21, 22). Under stress, the accumulation of MEcPP triggers a retrograde signalling cascade that facilitates calcium-dependent transcriptional reprogramming through the CALMODULIN-BINDING TRANSCRIPTION ACTIVATOR 3 (CAMTA3) transcription factor, and regulates adaptive growth by realigning auxin homeostasis and transport (23-25). While the role of chloroplast-derived metabolic signals is increasingly well-defined in photosynthesizing plants, the contribution of plastid metabolism to the progression of etiolated development remains largely unexplored (19, 26).

In recent decades, genome-wide association studies (GWAS) leveraging diverse natural populations have emerged as powerful tools for identifying genetic loci that regulate salinity stress responsiveness (27-31). In *Arabidopsis*, high-throughput phenotyping of shoot traits identified the protein kinase *salt-affected photosynthetic efficiency gene* (*SAPE*), a negative regulator of photosynthetic efficiency and growth maintenance during the early osmotic phase (32). Similarly, natural variation in the sodium transporter *HIGH AFFINITY K*^*+*^ *TRANSPORTER1 (HKT1)* demonstrates how *cis*-regulatory polymorphisms can drive adaptation; “weak” *HKT1* alleles facilitate tolerance in coastal populations by controlling Na^+^ translocation (27). Furthermore, variation at the *HKT1* locus influences salt-induced changes in root architecture, illustrating how natural regulatory diversity can shape complex, stress-adaptive developmental responses (29). GWAS in cereal crops has similarly linked natural variation in Na^+^ transport to salt tolerance. In rice, OsWRKY53 acts as a negative regulator of salt tolerance by directly repressing *OsHKT1;5* (33). Similarly, in maize, a naturally occurring rare variant of the plasma membrane antiporter *ZmSOS1* with a 4-bp frame-shifting deletion significantly impacts root Na^+^ efflux and shoot accumulation (34). Together, these examples demonstrate that natural variation in ion transport pathways is a fundamental mechanism utilized across diverse species to adapt to saline environments.

Here, we investigate the genetic basis of natural variation in salt-responsive apical hook development using the *Arabidopsis* HapMap population (35). Through an unbiased genome-wide association approach, we identify a chromosome 5 locus associated with salt sensitivity of hook bending and uncover regulatory variation within the promoter of *1-DEOXY-D-XYLULOSE 5-PHOSPHATE REDUCTOISOMERASE* (DXR), a key enzyme of the plastidial MEP pathway. These findings implicate plastid metabolism regulation as a previously unrecognized regulatory component of salt stress-modulated skotomorphogenesis.

## Materials and Methods

### Plant materials and growth conditions

274 *Arabidopsis thaliana* accessions of the HapMap population (35)(Table 1) were used for the GWAS analysis, where 5 seeds for each genotype were sown per treatment. The *pif1-2* mutant (Columbia-0 (Col-0) background) and *35S::TAP-PIF1* transgenic plants were described previously (36, 37). For the GWAS, and follow up haplotype phenotype analyses, seeds were sterilized using chlorine gas for three hours, followed by airing out for a subsequent 30 minutes. For all other assays, including phenotype screening, expression analyses by q-PCR, and protein isolation and immunoblotting, seeds were surface-sterilized in 0.05% Triton X-100 for 5 min, then in 1:1 household bleach:0.05% Triton X-100 for 10 min, and rinsed four times with sterile water. Sterilized seeds were sown on 50 µm nylon mesh (Sefar BV) over half-strength MS medium (Duchefa) with 0.8% plant agar (Duchefa), pH 5.7. After 4 d stratification at 4 °C in darkness, seeds were exposed to ∼125 μmol m^−2^ s^−1^ white light at 21 °C for 1 h, then wrapped in foil and grown vertically in darkness at 21 °C for 23 h. Seedlings were subsequently transferred under safe green light to new 0.5× MS plates with or without treatments and grown in darkness for 48 h unless otherwise stated. For salt stress experiments, 75 mM of NaCl (Fluka), was added to 0.5 MS media prior to autoclaving. A 10 mM stock solution of Fosmidomycin (FSM, Sigma) was made by dissolving directly in sterile water. This FSM stock solution was then added directly to warm media, to achieve a working concentration of 10 µM.

### GWAS Analysis

Variant data for the 274 accessions were obtained from the imputed 3M SNP dataset derived from the 1,001G panel (35). Variants were filtered to retain only those with a minor allele frequency greater than 0.05. Genome-wide association analyses were performed using GEMMA v0.98 (38), applying a univariate linear mixed model that accounts for population structure through a centered kinship matrix computed within GEMMA. Linkage disequilibrium (LD) was calculated using PLINK by estimating pairwise r^2^ values between the lead SNP and all variants within a ∼100Kb window (±50 Kb) (39).

### Gene Expression Analyses

Etiolated seedlings were harvested under dark conditions using a green safe light. ∼45 seedlings were collected per sample in 2ml safelock Eppendorf tubes containing two 1/8” steel ball bearings (Weldtite, 3906141), and were flash frozen in liquid nitrogen, before storing at -80 °C until extraction. Samples were ground twice at 25 Hz for 30 s using a Retsch shaker. RNA isolation was carried out using a EUR_X_ universal RNA kit (roboklon) in accordance with manufacturer instructions, with on column DNAse treatment (New England Biolabals) for 10 minutes. Reverse transcription was performed using the iScript− cDNA Synthesis Kit (BIO-RAD), with 1000 ng of RNA, and cDNA was diluted 1:10 with milliQ water. RT–qPCR was performed using a CFX Opus 384 Real-Time PCR System (Bio-Rad) with an initial denaturation at 95 °C for 3 min, followed by 40 cycles of 95 °C for 10 s and 60 °C for 30 s, with fluorescence acquisition at each cycle, followed by a melt curve analysis (55–95 °C, 0.5 °C increments). The iQ− SYBR® Green Supermix (Biorad) was used according to the manufacturers protocol with a final reaction volume of 5 µl. Relative gene expression was calculated as 2^–ΔΔCt with *ACTIN2* as the reference gene and normalized to Col-0 expression levels under control conditions. Primer sequences used for RT–qPCR were as follows: *DXR* (AT5G62790), forward 5′-TGAGGTTGCCCGACATCCTGAAGC-3′ and reverse 5′-AAGCACGAAAGGACCACCTGCG-3′; *ACTIN2* (AT3G18780), forward 5′-CTGGATCGGTGGTTCCATTC-3′ and reverse 5′-CCTGGACCTGCCTCATCATAC-3′. For expression of QTL5 targets and *PIF1* across different seedling organs under 75 mM NaCl and control, normalized count data were obtained from RNA-sequencing data of a previously performed experiment (18).

### Apical Hook Phenotyping

Photos of seedlings were taken with a Canon EOS R10 camera two days after transfer to treatment media (three days old in total). Phenotype analyses were carried out in Fiji (40), whereby apical hook angle was measured using the angle tool as described previously (41). For determining apical hook ratio, accessions with a mean hook angle of less than 90 ° under control conditions were omitted.

### Protein isolation and immunoblotting

Three-day-old *35S::TAP-PIF1, pif1-*2, and Col-0 seedlings were harvested in the dark under a green safe-light. Forty etiolated seedlings per sample were flash-frozen in liquid nitrogen and stored at – 80 °C in 2 mL safelock tubes with two 1/8” steel ball bearings. Samples were ground twice at 25 Hz for 30s, resuspended in 80 uL protein extraction buffer (50 mM Tris pH 7.5, 200 mM NaCl, 4 M urea, 0.1% Triton X-100, 5 mM DTT, cOmplete protease inhibitor cocktail tablet (Roche)), followed by centrifuging at 13,000 rpm at 4 °C for 15 min. Supernatants were mixed with 5 x laemmli buffer (300 mM Tris-HCl pH 6.8, 5% SDS, 50% glycerol, 100 mM DTT, 0.05% bromophenol blue), heated to 95 °C for 5 mins and kept on ice. Proteins were separated on 10% SDS-PAGE gels and transferred to nitrocellulose membranes using the Trans-Blot Turbo system (Bio-Rad). Membranes were blocked in 5% milk/TBST for 1 h, then incubated overnight at 4 °C with primary antibodies: anti-Myc (1:1000, Santa Cruz 9E10 sc-40), or anti-actin (1:1000 Santa Cruz sc-47778). After washing, membranes were incubated with anti-mouse (1:2000 PhytoAB PHY6006) secondary antibody for 1 h at room temperature, washed again, and visualized with Clarity ECL substrate on a ChemiDoc system (Bio-Rad). Signal intensity of protein bands was quantified using Image Lab software, volume tool (Bio-Rad, version 6.1).

### Statistical analysis and data representation

Statistical analyses were carried out in RStudio (version 4.4.1). Information regarding used statistical methods are provided in the figure legends. All plots were generated using the R package ggplot2, and haplotype heatmaps were made using the R package pheatmap.

## Results

### Natural Variation in Apical Hook Salt Sensitivity is Driven by Growth Under Salinity

Our previous work showed that salinity reduces apical hook bending in etiolated *Arabidopsis thaliana* Col-0 seedlings (18). To determine whether salt responsiveness of apical hook growth varies among natural accessions and to identify potential genetic bases, we quantified salt responses across the Arabidopsis HapMap population and performed a genome-wide association study. Salt responsiveness was measured as the ratio of apical hook angle of three day old seedlings exposed to 75 mM NaCl relative to control conditions (0.5 MS media). Substantial variation in apical hook angle was observed across accessions under both control and salt stress conditions, as well as in the ratio between the two (Fig. 1A). All but one accession had an apical hook ratio below 1, highlighting that salt has an inhibitory effect on apical hook bending across accessions. Although apical hook angle under salt stress correlated with baseline hook angle, salt responsiveness was only weakly related to control growth and instead was strongly determined by hook angle under NaCl treatment (Fig. S1). These results indicate that variation in apical hook ratio is driven primarily by growth behaviour under NaCl rather than by baseline apical hook curvature.

**Figure 1.**
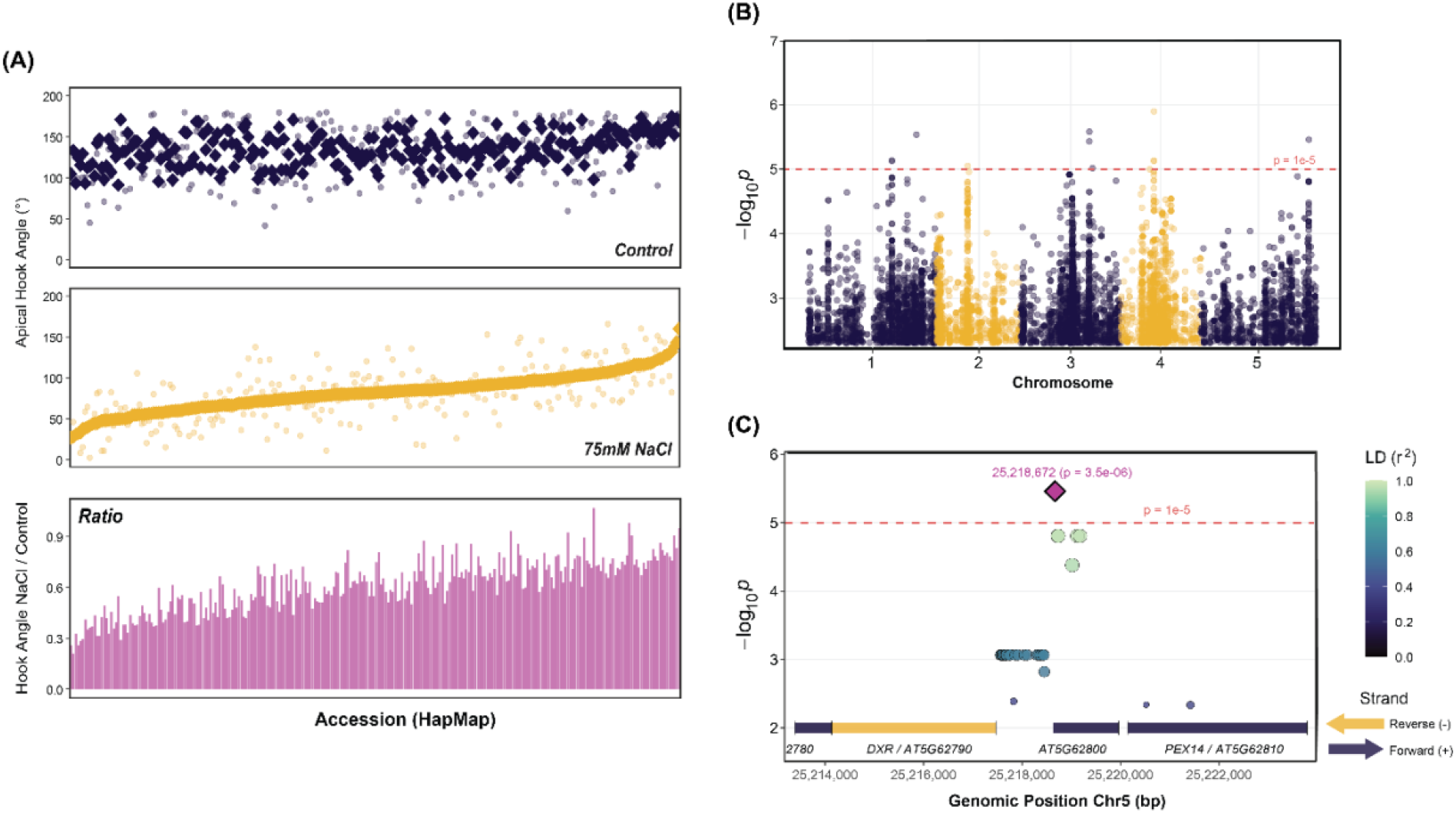
Genome Wide Association Study (GWAS) based on ratio between apical hook angle in control conditions / apical hook angle under 75 mM NaCl. (A) Natural variation in apical hook curvature across the HapMap population (274 accessions). Mean apical hook angles under control (navy; top) conditions and 75 mM NaCl (gold; middle), where diamonds represent means, and dots represent individual measurements. Apical hook ratio (pink; bottom) calculated by dividing mean angle under NaCl by mean angle under control conditions for each accession. Physiology data ordered from smallest to largest mean hook angle under 75 mM NaCl. (B) Manhattan plot representing output of genome-wide association analysis of salt responsiveness (apical hook angle ratio). Dashed line indicates the genome-wide significance threshold of p=1×10^-^5,. (C) Regional association plot of the chromosome 5 QTL surrounding the lead SNP at position 25,218,673 (p = 3.5×10^−6^, pink diamond). SNPs are coloured by linkage disequilibrium (r^2^) with the lead variant. Gene boundaries are shown below, with arrows indicating strand / transcriptional direction.

### A chromosome 5 QTL is associated with variation in salt-induced reduction of apical hook bending

After excluding non-germinating accessions or those with impaired apical hook formation under control conditions (<90°), 274 accessions were retained for genome-wide association analysis (Table 1). Association mapping of salt responsiveness identified multiple putative loci across all five chromosomes (Fig. 1B). When considering association peaks with a high minor allele frequency (> 0.3), two prominent signals were detected on chromosomes 1 and 5, with lead SNPs at positions 19,929,998 (−log_10_(*p*) = 5.13) and 25,218,672 (−log_10_(*p*) = 5.46) respectively. We focused subsequent analyses on the chromosome 5 locus, prioritizing genes within a region of moderate linkage disequilibrium (r^2^ ≥ 0.55) surrounding the lead SNP, which includes several functionally annotated genes. (Fig. 1C, Fig. S2). Within this region four candidate genes appeared ranging from *AT5G26790* to *AT5G26820* (Fig. S2).

The lead SNP on chr5 was located withing the CDS of *AT5G62800*, 38bp from the transcriptional start site (TSS). Whilst this gene provides an interesting target for follow, formerly performed RNA-sequencing of etiolated seedlings grown under salt stress or control conditions reveal no reads of *AT5G62800* across cotyledons, apical hooks, or hypocotyls (Fig. S2B) (18). We did however observe expression for *AT5G26790, AT5G267810*, and *AT5G67820* (Fig. S2B) The next gene with closest proximity to the lead SNP is *AT5G26790* encoding *1-DEOXY-D-XYLULOSE 5-PHOSPHATE REDUCTOISOMERASE* (*DXR*). RNA-sequencing showed expression in all tested seedling organs, with the highest counts in the cotyledons. Interestingly, we observed pronounced natural variation within the shared promoter region of *AT5G26790* and *AT5G62800*. This region segregates into two major haplotypes across the HapMap population: one corresponding to the reference Col-0–like promoter sequence and a second defined by a set of alternative SNPs that are inherited together (Fig. S3A). The alternative promoter haplotype was associated with reduced salt responsiveness of apical hook growth relative to the reference haplotype (Fig. S3C). These promoter variants are in moderate linkage disequilibrium with the lead SNP at the chromosome 5 locus (r^2^ ≈ 0.62), suggesting transcriptional regulation of either gene could confer reduced salt responsiveness. Considering that *DXR* is known to be transcriptionally repressed by the PIFs in etiolated seedlings (42), and has been found to be implicated in salt stress responses (43, 44), as well as hormone biosynthesis (45) we selected *DXR* as a candidate for follow up analyses.

### Differences in salt response correlate to two major *pDXR* haplotype variants

Analysis of SNP variation within the shared promoter of *DXR* and *AT5G62800* revealed two major haplotypes that segregate among the HapMap accessions, with the alternative haplotype associated with reduced salt sensitivity (Fig. S3C). To independently validate this association, we selected 15 accessions representing the two haplotype groups and repeated the apical hook phenotyping assay with increased sample size (Fig. 3A-B). Under control conditions, accessions carrying the alternative *pDXR* haplotype exhibited modestly increased apical hook angles relative to those with the reference haplotype (Fig. 3B, 3C). Notably, the magnitude of salt induced reduction in apical hook bending differed between haplotypes, with accessions carrying the alternative haplotype maintaining significantly higher apical hook angles under 75 mM NaCl (Fig. 3B, 3C). Consistent with this observation, a significant haplotype × treatment interaction was detected (p = 2.87 × 10^−6^), indicating that the effect of haplotype on apical hook angle differs between control and salt stress conditions.

**Figure 2.**
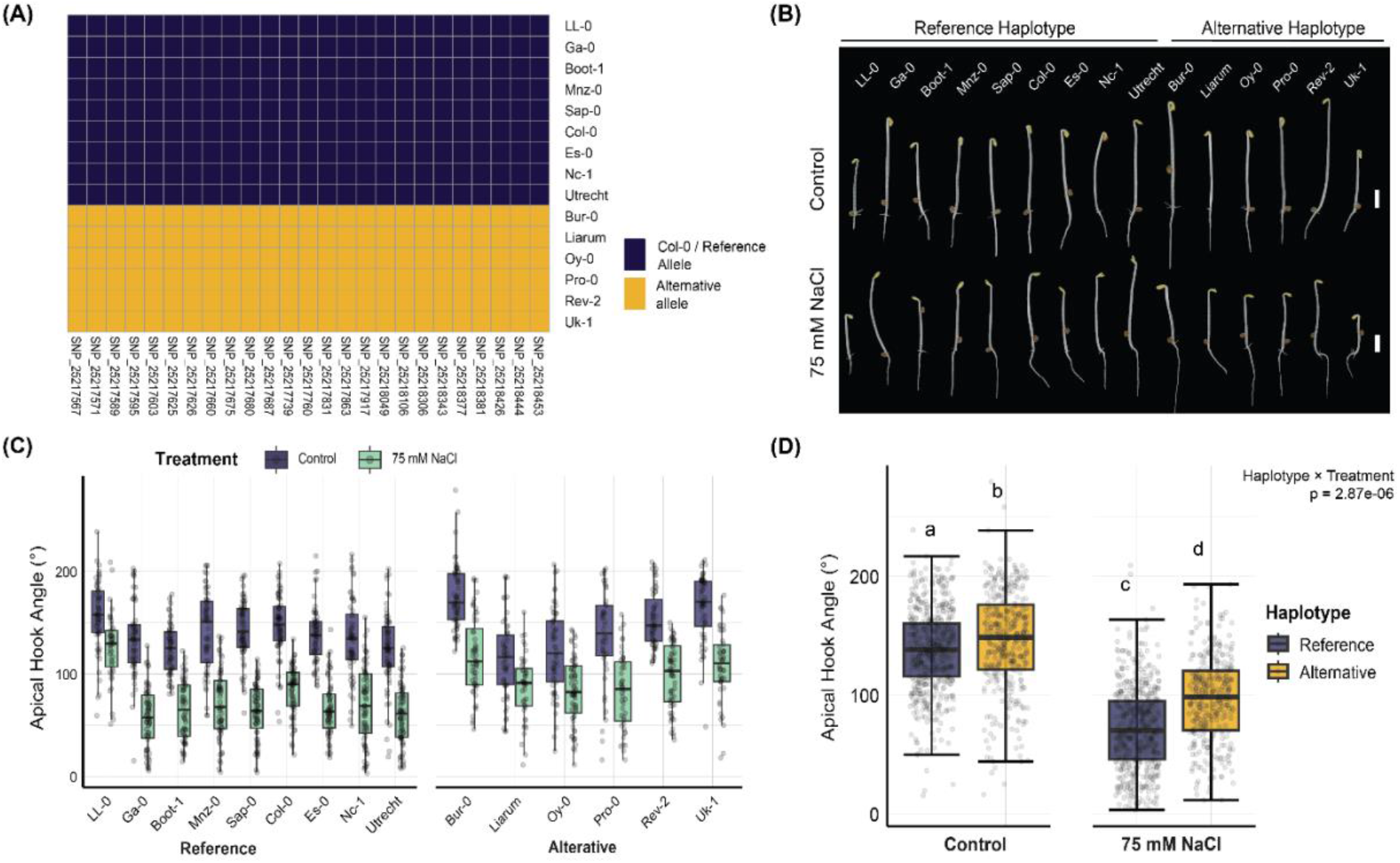
Independent validation of association between alternative haplotype variant and apical hook angle under salt stress. (A) Haplotype matrix displaying allelic variation across SNPs within the shared promoter of *DXR* and *AT5G62800* for 15 *Arabidopsis* accessions. (B) Representative phenotype of seedlings belonging to the reference (left) or the alternative (right) haplotype groups grown under 75 mM NaCl and control conditions. Scale bar = 2 mm. (C) Boxplot representing apical hook angles under 75 mM NaCl and control conditions in accessions of the reference haplotype group (left) or of the alternative haplotype group (right). (C) Boxplot representing apical hook angles under 75 mM NaCl and control conditions in accessions (grouped) of the reference haplotype group or of the alternative haplotype group. Significance of interaction effect between haplotype group and treatment calculated by two-way ANOVA, and significance letters denote statistically significant differences (Tukey’s post hoc test, *p* < 0.05). Boxplots display median, interquartile range, and whiskers extending to 1.5× IQR.

**Figure 3.**
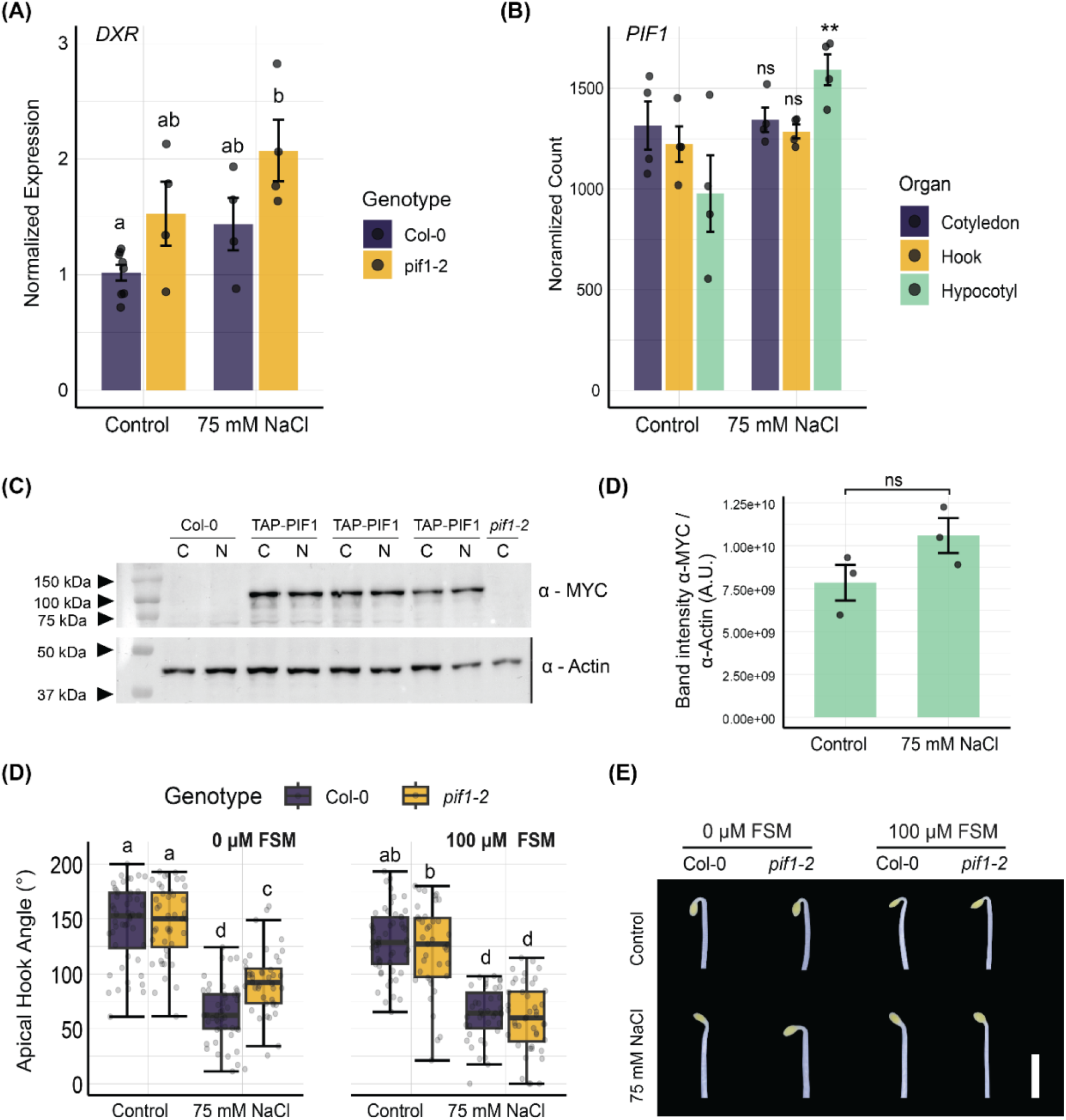
Loss of PIF1 Increases Hook Bending Under NaCl via Increased MEP Pathway Activity. (A) Relative *DXR* transcript levels measured by RT–qPCR in etiolated Col-0 and *pif1-2* seedlings grown under control or 75 mM NaCl conditions. Expression was normalized to *ACTIN2* and normalized to Col-0 control. Bars represent mean ± SE; dots indicate biological replicates. Different letters denote statistically significant differences (two-way ANOVA with Tukey’s post hoc test, *p* < 0.05). (B) Normalized *PIF1* transcript abundance across cotyledons, apical hooks, and hypocotyls under control and salt stress conditions, derived from RNA-sequencing data. Statistical significance calculated using DESeq2 pairwise comparison per organ and is indicated (\*\**p* < 0.01; ns, not significant). (C) Immunoblot analysis of 35S promoter driven TAP-PIF1 protein levels in etiolated seedlings grown under control (C) or 75 mM NaCl (N) conditions. Bar plot shows the PIF1-TAP/actin ratio of band intensity. C=(D) Apical hook angles of Col-0 and *pif1-2* seedlings grown under control or 75 mM NaCl conditions in the absence or presence of 100 μM fosmidomycin (FSM). Different letters denote statistically significant differences (three-way ANOVA with Tukey’s post hoc test, *p* < 0.05). Barplots display mean and ± standard error. Boxplots display median, interquartile range, and whiskers extending to 1.5× IQR. (E) Representative apical hook phenotype of Col-0 and *pif1-2* seedlings grown under 75 mM NaCl and control conditions, either with or without a 100 μM fosmidomycin. Scale bar = 2 mm.

### PIF1 partially regulates apical hook sensitivity to NaCl via the MEP pathway

The identification of *DXR* as a candidate gene underlying natural variation in salt-responsive apical hook bending prompted us to examine how altered *DXR* expression levels affect etiolated growth. Since the transcription of *DXR* in the dark is known to be repressed by a major regulator of seedling etiolation, transcription factor PHYTOCHROME INTERACTING FACTOR1 (PIF1) in the dark (42), we examined the expression of *DXR* under salt stress and control conditions, in the *pif1-2* mutant and Col-0. Both salt treatment and loss of PIF1 increased *DXR* transcript levels, with the highest expression observed in salt-treated *pif1-2* seedlings, significantly differing from wild-type seedlings grown under control (Fig. 3A). The absence of a significant two-way ANOVA genotype × treatment interaction suggests that NaCl and PIF1 regulate *DXR* transcription additively in the Col-0 background. Consistent with this, *PIF1* transcript levels were largely unchanged by salt treatment in relevant tissues (cotyledons and apical hooks)(Fig. 3B). Moreover, immunoblot analysis revealed non-significant TAP-PIF1 protein levels across treatments (Fig. 3C). Together, these results show that both salt stress and PIF1 contribute to the regulation of *DXR* expression during etiolation.

Next, we tested whether increased *DXR* expression in *pif1-2* seedlings affects the apical hook salt response. We observed significantly higher apical hook angles in salt grown *pif1-2* mutants when compared to Col-0 (Fig. 2A). To determine whether this enhanced hook bending was dependent on elevated *DXR* activity, we inhibited DXR enzymatic function using fosmidomycin (FSM), which blocks the DXR-catalysed conversion of 1-deoxy-D-xylulose 5-phosphate (DXP) to MEP, thereby acutely reducing flux through the plastidial MEP pathway (46). Indeed, FSM treatment reduced *pif1-2* apical hook angles to the Col-0 level under 75 mM NaCl, indicating that the increased apical hook angle under salinity in *pif1-2* is caused by increased flux through the MEP pathway, consistent with increased *DXR* expression in the absence of PIF1.

### *DXR* expression is correlated with apical hook angle under NaCl in natural *Arabidopsis* accessions

We have shown that variation in *DXR* promoter sequences, and *DXR* transcript levels are associated with differences in salt sensitivity in terms of apical hook curvature. To integrate these observations, we selected eight previously phenotyped accessions and tested whether *DXR* expression differs between the two promoter haplotype groups under salt stress and control conditions. *DXR* expression did not significantly differ between haplotype groups under control conditions, but was significantly elevated in the alternate haplotype under salt stress (Fig. 4A), confirming a stress-dependent effect of promoter variation on gene expression. To further examine the relationship between *DXR* expression and apical hook growth, we compared mean *DXR* transcript levels with mean apical hook angle across these accessions (and *pif1-2*) under control and salt stress conditions. Apical hook angle did not significantly correlate with *DXR* expression under control conditions (r = 0.284, p = 0.459), whereas a strong positive correlation was observed under NaCl treatment (r = 0.704, p = 0.034) (Fig. 3B). Consistently, salt responsiveness, measured as the NaCl/control ratio of apical hook bending, positively correlated with *DXR* expression under salt stress (r = 0.772, p = 0.015) (Fig. 4C). Together, these results link natural variation in *DXR* expression to salt-responsive apical hook growth and support a role for *DXR* and the MEP metabolic pathway in skotomorphogenic development under salinity.

**Figure 4.**
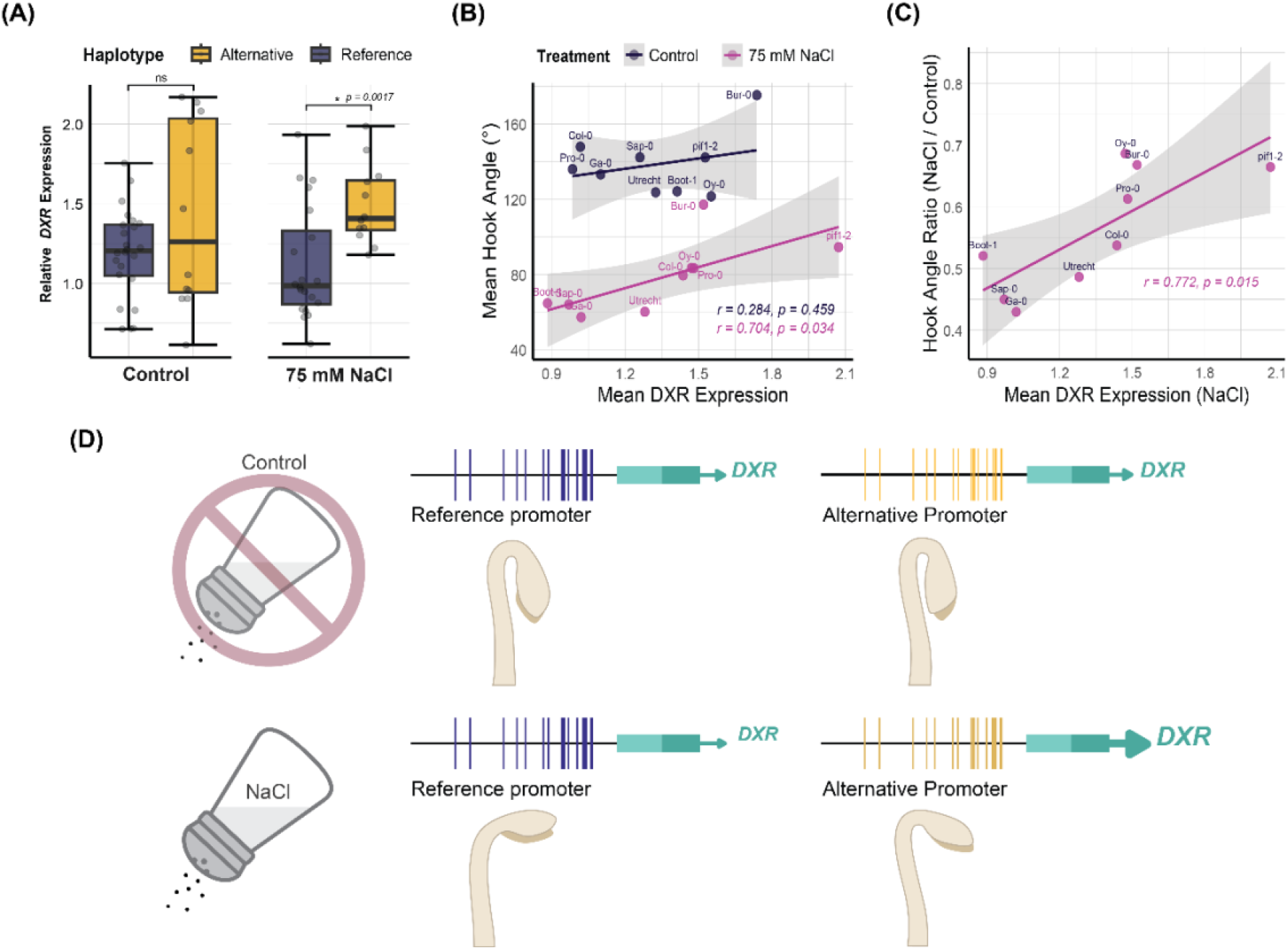
Natural variation in *DXR* expression under salt stress correlates with apical hook angle ratio (NaCl/Control). (A) Relative *DXR* expression in accessions carrying the reference or alternative *DXR* promoter haplotypes under control and 75 mM NaCl conditions. Statistical significance determined by t-test (\**p* < 0.05). (B) Mean apical hook angle plotted against mean *DXR* expression for selected natural accessions and *pif1-2* under control and 75 mM NaCl conditions. Linear regressions are shown with shaded 95% confidence intervals. (C) Apical hook angle ratio (NaCl/Control) plotted against mean *DXR* expression measured under salt stress. Linear regression and 95% confidence interval are shown. Boxplots display median, interquartile range, and whiskers extending to 1.5× IQR. (D) Schematic of *DXR* promoter haplotypes showing that the alternative haplotype confers increased *DXR* expression and enhanced apical hook bending under salt stress, but not under control conditions.

## Discussion

Salt stress has distinct impacts on skotomorphogenesis, restricting hypocotyl elongation and disrupting apical hook formation (18, 47). That being said, the genetic basis underlying natural variation in salt stress-responsive physiological changes remains poorly understood. GWAS analysis has previously been implemented to identify genetic components that drive natural variation responses to salinity, including root system architecture remodelling, photosynthetic efficiency changes, and root gravitropic responses (29, 32, 48). Here, we combined GWAS with molecular and physiological analyses to identify *DXR*, a key enzyme of the plastidial MEP pathway (49), as a putative regulator of salt-responsive apical hook bending in etiolated *Arabidopsis* seedlings.

Phenotypic screening of natural *Arabidopsis* ecotypes of the HapMap population revealed that salinity broadly supresses hook bending. These findings are consistent with our previous data pertaining to Col-0, but here accessions displayed variability in the extent of this inhibition (Fig. 1A). Notably, salt sensitivity in terms of apical hook correlates strongly with apical hook angle under salt stress, but only weakly with the angle under control (Fig. S1). This suggests that salt stress actively reshapes developmental programs rather than scaling pre-existing growth differences. Genome-wide association analysis identified a putative quantitative trait locus on chromosome 5, and calculating linkage disequilibrium with the lead SNP presented a narrow interval containing few candidate genes. Although the lead SNP was located within the coding region of *AT5G62800*, these SNPs do not induce non-synonymous mutations (Table 1). In addition, the lack of detectable expression in etiolated seedlings under our experimental conditions argued against its involvement. In contrast, the adjacent gene *DXR* emerged as a compelling candidate based on the presence of substantial genetic variation within its shared promoter region with *AT5G62800*, and it’s abundant expression in the cotyledons and apical hooks of etiolated seedlings (Fig S2B).

Interestingly, the *DXR* promoter region segregates into two major haplotypes across the HapMap population, and the alternative haplotype associates with reduced salt sensitivity of apical hook bending (Fig. S3A & C; Fig. 4D). Independent phenotypic validation across selected accessions confirmed that this haplotype confers enhanced maintenance of hook curvature specifically under salt stress (Fig. 2), supporting the theory that regulatory variation at this locus modulates stress-induced growth responses. In most land plants, including *Arabidopsis*, and algae the *DXR* gene is encoded as a highly evolutionarily conserved, single-copy enzyme, catalysing the first committed step of the plastid-localized MEP pathway (50, 51). With consideration for this conservation, it is logical that genetic variation would be found in the promoter sequence of *DXR*, which contains the essential cis-regulatory elements required for spatial, temporal, and environmental control of the gene (42, 50). Promoter variation may fine-tune *DXR* transcription and MEP pathway flux to suit the metabolic demands of the local environment. This regulatory mechanism is critical because MEP-derived isoprenoids serve as indispensable building blocks for an array of hormones and active metabolites, including chlorophylls, carotenoids, gibberellins, and abscisic acid, which are fundamental for tuning plant physiology in response to the environment (42, 50-55).

The transcriptional regulation of *DXR* was previously shown to be governed by master light-regulated transcription factors such as ELONGATED HYPOCOTYL 5 (HY5) and the PIFs, which regulate MEP pathway flux under dark or light conditions (42). Since the direct transcriptional repression of *DXR* by PIF1 in the dark has been experimentally verified (42), we investigated the combined effects of salinity and loss of PIF1 in hook bending. Our results indicate that salt stress and the *pif1-2* mutation independently and additively increase *DXR* expression, which is associated with increased hook bending under salt stress but not control conditions (Fig. 3 A & D). The reversal of the enhanced apical hook bending in salt-treated *pif1-2* seedlings upon fosmidomycin treatment supports a requirement for increased flux through the MEP for reduced salt sensitivity of *pif1-2*. This relationship was further supported by natural variation analyses, which revealed a strong positive correlation between *DXR* expression and apical hook bending specifically under salt stress, but not under control conditions (Fig. 4 B & C). Notably, in contrast to a previous report proposing salt-induced reductions in PIF1 abundance (56) we did not observe significant changes in PIF1 protein levels in etiolated seedlings following salt treatment.

The precise mechanisms by which altered *DXR* expression under salinity modulates skotomorphogenic development remains to be determined. Nevertheless, we present a plausible pathway that merits consideration. Recent work has proposed the MEP pathway as an oxidative stress sensing and response system, largely due to the sensitivity of its terminal iron–sulphur cluster enzymes, HDS (4-hydroxy-3-methylbut-2-enyl diphosphate synthase / IspG) and HDR ( 4-hydroxy-3-methylbut-2-enyl diphosphate reductase / IspH), to reactive oxygen species (ROS) (19, 22). Since salt stress is a known inducer of ROS accumulation (57), oxidative inhibition of HDS could create a metabolic bottleneck within the pathway, leading to accumulation of the intermediate metabolite 2-C-methyl-D-erythritol-2,4-cyclodiphosphate (MEcPP) (21). MEcPP is well established as a plastid-derived retrograde signalling molecule involved in both biotic and abiotic stress responses (21, 54), presenting it as a compelling candidate mediator of salt-responsive apical hook opening. Under salt stress conditions, transcriptional upregulation of *DXR* would be expected to further enhance flux into the MEP pathway, thereby amplifying MEcPP accumulation. This model provides a potential explanation for why *DXR* expression correlates with apical hook curvature specifically under salt stress, but not under control conditions (Fig. 4).

MEcPP has previously been shown to restrict plant growth by modulating auxin biosynthesis and transport, including reductions in PIN1 protein abundance, resulting in shortened hypocotyls (23, 25, 58). At first glance, this appears at odds with our earlier observation that salt stress reduces auxin signalling in the apical hook of Col-0 seedlings (18). However, a key distinction is that MEcPP-mediated regulation of auxin responses has been shown to be light dependent (58, 59). In dark-grown seedlings, auxin levels remain largely comparable across genotypes with differing MEcPP status (58), suggesting that MEcPP signalling may exert context-dependent effects on growth. We therefore propose that elevated *DXR* expression and MEP pathway activity under salt stress could contribute to stress-dependent modulation of apical hook development during etiolation. A key challenge for future studies will be to uncover how plastid-derived metabolic and signalling outputs intersect with hormonal and growth-regulatory pathways to shape stress-responsive development, particularly in the dark.

## Abbreviations

GWAS: Genome wide association study
MEP: methylerythritol phosphate
DXR: 1-DEOXY-D-XYLULOSE 5-PHOSPHATE REDUCTOISOMERASE
PIF1: Phytochrome interacting factor1

## Supplementary data

Table S1. List of *Arabidopsis thaliana* ecotypes used for GWAS analysis, with the average hook ratio (NaCl/control).

Figure S1. Salt responsiveness is predominantly driven by the salt-treated phenotype rather than the control phenotype.

Figure S2. Candidate genes underlying a GWAS locus on Arabidopsis chromosome 5.

Figure S3. Alternative chromosome 5 Lead SNP and *pDXR* haplotypes associated with reduced salt sensitivity.

Table S2. Raw GWAS output.

## Acknowledgements

We thank Kyra van der Velde and Ronald Pierik for providing the *pif1-2* mutant, and Gabriela Toledo-Ortiz for the *35S::TAP-PIF1* transgenic *Arabdiopsis thaliana* lines. We are grateful to Leo Willems for generating and providing the *Arabidopsis* HapMap seed collection used for the GWAS. We also extend thanks to Cornelia Rommens for assistance with haplotype follow-up studies.

## Author contributions

E.v.V. and C.M.M.G. planned and designed the research; E.v.V., T.v.d.D. and X.C. performed experiments, E.v.V., R.B. and J.A.D. performed data analysis, E.v.V. and C.M.M.G. wrote the manuscript, all authors provided feedback on the manuscript.

---

**Figure S1.**
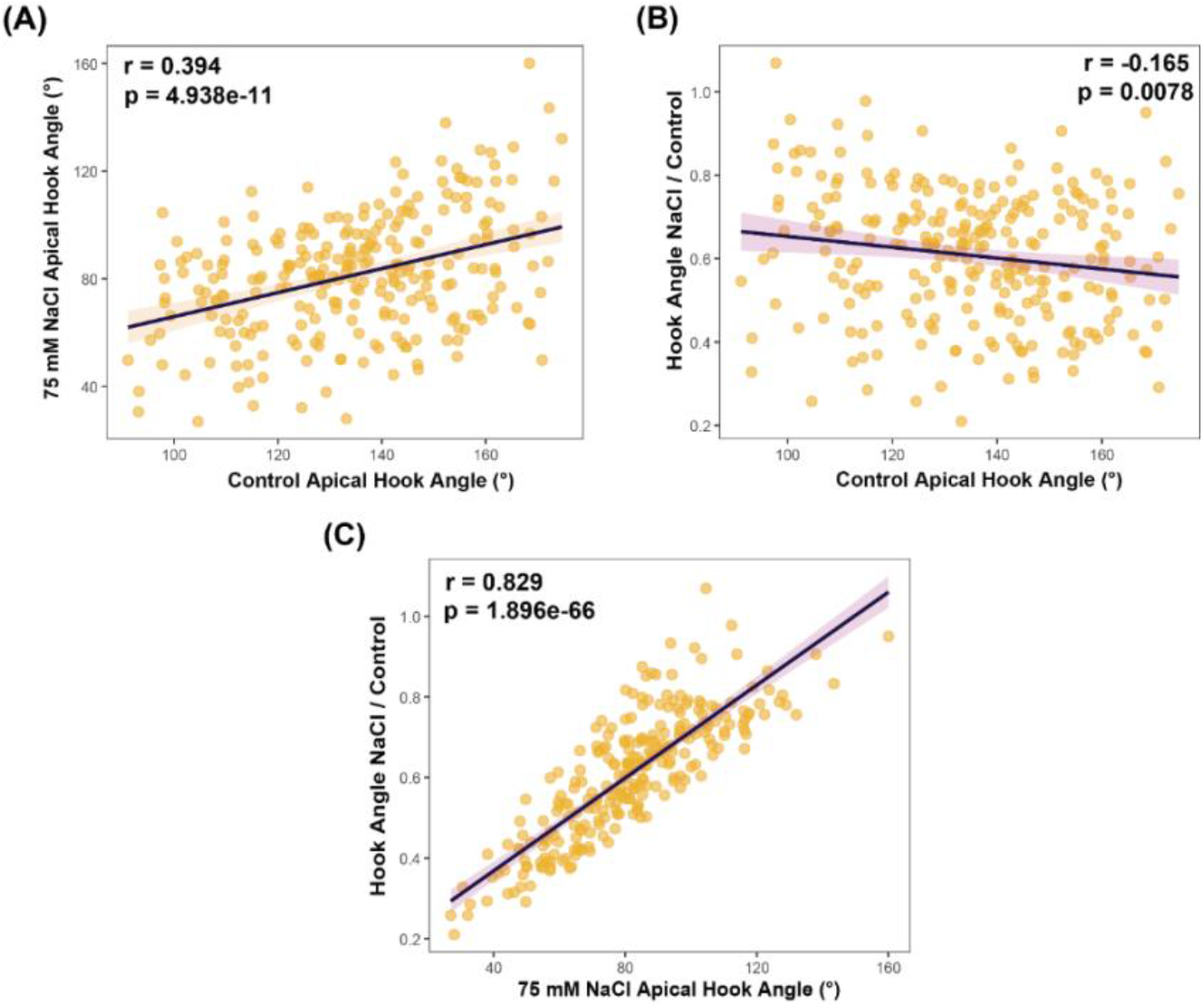
Salt responsiveness is predominantly driven by the salt-treated phenotype rather than the control phenotype. Correlations among apical hook angle phenotypes across Arabidopsis accessions as follows. (A) Apical hook angle under control conditions plotted against hook angle under 75 mM NaCl treatment (r = 0.394, *p* = 4.94×10^−11^). (B) Hook angle ratio (NaCl/Control) plotted as a function of apical hook angle under control conditions (r = −0.165, *p* = 0.0078). (C) Hook angle ratio (NaCl/Control) plotted against apical hook angle under 75 mM NaCl treatment (r = 0.829, p = 1.90×10^−66^). Lines represent linear regression fits with 95% confidence intervals (shaded regions). Each point represents an individual HapMap accession.

**Figure S2.**
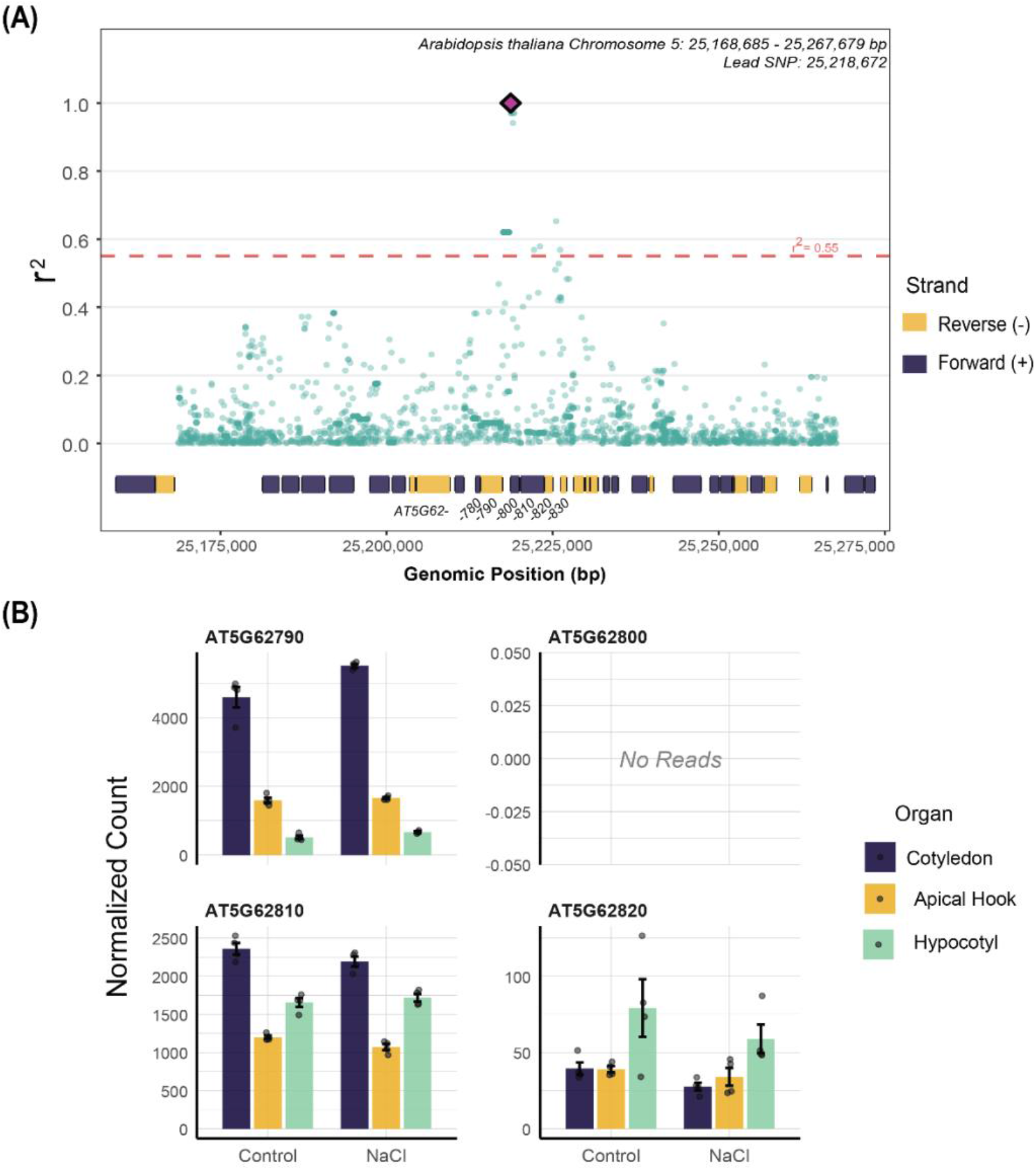
Candidate genes underlying a GWAS locus on Arabidopsis chromosome 5. (A) Linkage disequilibrium (r^2^) plot showing SNPs in the region surrounding the lead SNP (diamond) at position 25,218,672 bp. The dashed red line indicates the r^2^ = 0.55 threshold. Gene models are shown below, with forward strand (+) genes in purple and reverse strand (−) genes in yellow. (B) Normalized expression counts from DESeq2 for four candidate genes (AT5G62790 / DXR, AT5G62800, AT5G62810 / PEX14, AT5G62820) in three seedling organs under control and NaCl (salt stress) conditions. Data represent mean ± SE (n = 4 biological replicates). AT5G62800 showed no detectable expression. None of the genes showed statistically significant differential expression in response to salt treatment after FDR correction.

**Figure S3.**
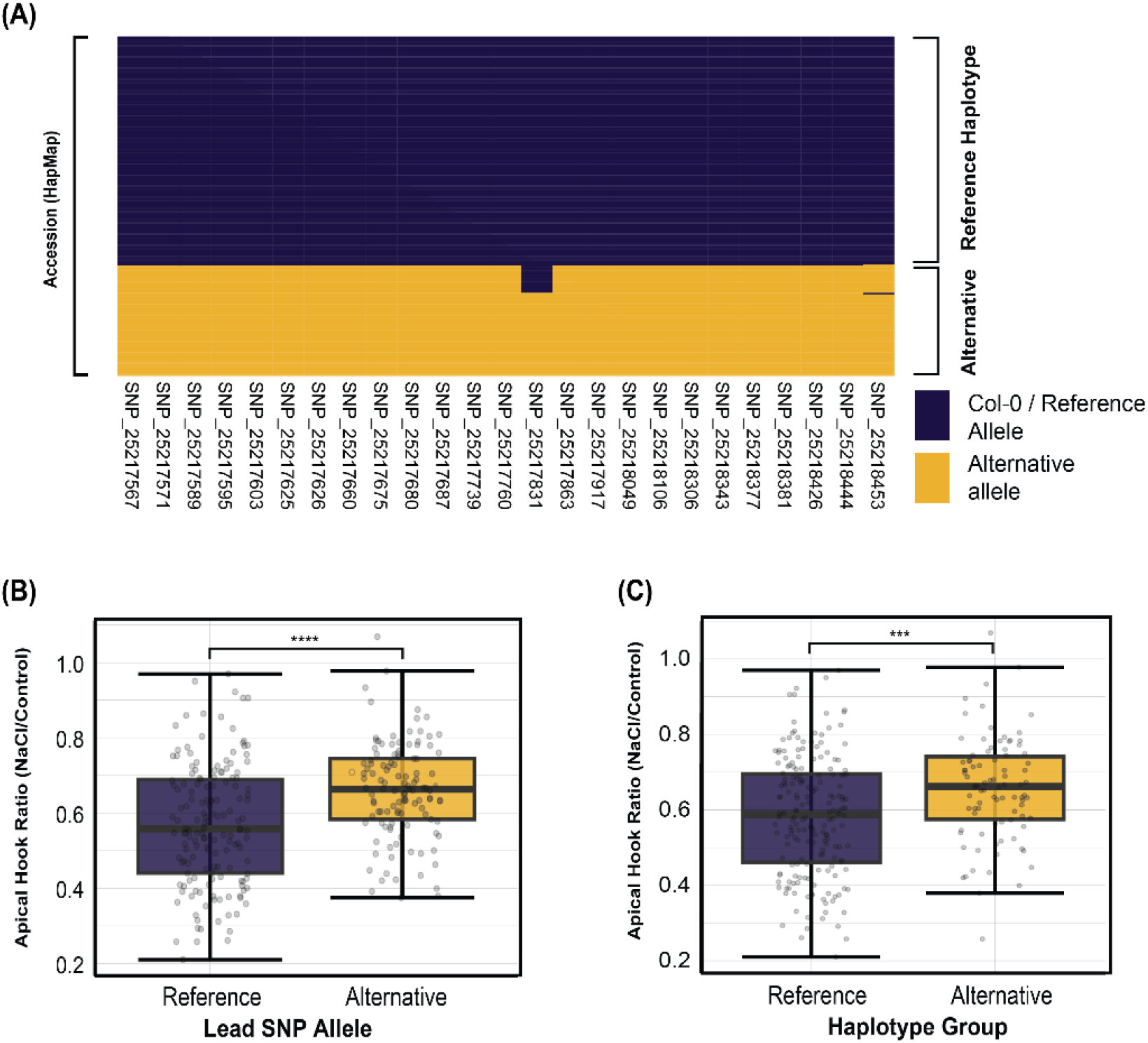
Alternative chromosome 5 Lead SNP and *pDXR* haplotypes associated with reduced salt sensitivity. (A) Haplotype matrix displaying allelic variation across SNPs within the shared promoter of *DXR* and *AT5G62800*. reference (Col-0) alleles shown in purple and alternative alleles in gold. (B) Boxplot representing difference in apical hook ratio in accessions with alternative lead SNP (gold) or the reference lead SNP (purple). (C) Boxplot representing difference in apical hook ratio in accessions of the alternative haplotype group (gold) or of the reference haplotype group (navy). Box plots display median, interquartile range, and whiskers extending to 1.5× IQR; individual accessions are shown as points. Statistical significance determined by t-test (\*\*\**p* < 0.001, \*\*\*\**p* < 0.0001).

